# Ischemic Stroke Reduces Bone Perfusion and Alters Osteovascular Structure

**DOI:** 10.1101/729632

**Authors:** Nicholas J. Hanne, Andrew J. Steward, Carla Geeroms, Elizabeth D. Easter, Hannah L. Thornburg, Greet Kerckhofs, Tatjana Parac-Vogt, Huaxin Sheng, Jacqueline H. Cole

## Abstract

**Rationale:** Stroke patients lose bone mass and experience fracture at an elevated rate. Although functional intraosseous vasculature is necessary for skeletal maintenance, the effect of stroke on osteovasculature is unknown.

**Objective:** To characterize changes to osteovascular function, structure, and composition following mild-to-moderate-severity ischemic stroke in mice, both with and without exercise therapy.

**Methods and Results:** Twelve-week-old male mice (n=27) received either a stroke (middle cerebral artery occlusion) or sham procedure, followed by four weeks of daily treadmill or sedentary activity. Intraosseous perfusion, measured weekly in the proximal tibial metaphysis, was reduced by stroke for two weeks. In the second week of recovery, exercise nearly restored perfusion to sham levels, and perfusion tended to be lower in the stroke-affected limb. At the conclusion of the study, osteovascular structure was assessed with contrast-enhanced computed tomography in the distal femoral metaphysis. Stroke significantly increased osteovascular volume and branching but reduced the relative number of blood vessels close to bone surfaces (6-22 μm away) and increased the relative number more than 52 μm away. These differences in vessel proximity to bone were driven by changes in the stroke-exercise group, indicating compounded effects of stroke and exercise. Exercise, but not stroke, nearly reduced the amount of osteogenic Type H blood vessels in the proximal tibial metaphysis, quantified with immunofluorescence microscopy.

**Conclusions:** This study is the first to examine the effects of stroke on osteovasculature. Stroke increased the amount of osteovasculature, but since blood vessels close to bone are associated with bone remodeling, the shift in osteovascular structure could play a role in bone loss following stroke. The exercise-induced reduction in the amount of Type H vessels and the stroke-exercise effect on osteovascular structure suggest moderate aerobic activity may have detrimental effects on bone remodeling during early stroke recovery.

## Introduction

Stroke is the leading cause of disability in the US, as well as one of the leading causes worldwide [1]. Stroke not only causes cognitive and motor impairments but also negatively affects skeletal health [2–8]. Stroke patients lose bone mass at an accelerated rate, fall more, and experience hip fracture 2-4 times more frequently than individuals of the same age who have not experienced stroke [2,3,5]. Because bone adapts to altered mechanical loading [9], and bone mineral density (BMD) loss and fracture preferentially affect the paretic limbs [6,7], hemiplegia and limb disuse are thought to be solely responsible for these skeletal deficits post-stroke [4,6,8]. In addition, rehabilitation activity and exercise are correlated with reduced fracture risk and improved skeletal health outcomes following stroke [3,10–13], further supporting loading as a primary driver. However, a study in severe stroke patients showed lower BMD in various sites on the stroke-affected side, relative to the unaffected side, despite complete bedrest [8]. Even after controlling for activity level and other factors, lower vascular elasticity, a metric of vascular health, has been associated with lower polar stress-strain index, a surrogate measure of bone torsional stiffness and strength [14], in the radial diaphysis [15]. Together these results suggest that the negative effects of stroke on bone extend beyond just mechanical unloading and that vascular deficits may also contribute.

Because vasculature is critically important for maintaining functional bone tissue, supplying not only oxygen and nutrients but also cell signaling factors [16], vascular dysfunction may contribute to the increased fracture risk experienced by stroke patients. Little is known about the effects of stroke on limb vasculature, in particular on vascular perfusion and structure, and no previous study has examined the effects on the vasculature within bone, or *osteovasculature.* Two previous studies reported decreased blood flow in the affected lower leg compared to the unaffected leg in elderly chronic stroke patients with mild-to-moderate gait asymmetry at least 6 months after moderate ischemic stroke [17,18]. Changes to limb blood flow, however, may not be indicative of changes to osteovasculature, which has more direct impacts on bone remodeling and the marrow microenvironment. In osteoporotic individuals, increased intraosseous perfusion measured by contrast-enhanced clinical imaging was strongly correlated with increased bone formation rate in the iliac crest [19] and weakly correlated with higher BMD in the proximal femur [20]. In addition, an increased number of capillaries have been observed within 50 μm of bone surfaces at active remodeling sites, compared to non-remodeling bone surfaces, in both humans [21] and rats [22]. Within long bones, endothelial cells that express both endomucin (EMCN) and CD31 *(Type H cells)* have been shown to regulate osteogenesis in mice [23,24]. The effects of stroke on these osteovascular metrics (perfusion, structure, and cellular composition) are unknown and may provide insight into the relationship between bone vasculature and bone loss following stroke. Although characterizing osteovasculature in humans is difficult, it can be examined in animal models, such as the established rodent models of ischemic stroke involving middle cerebral artery occlusion (MCAo) [25,26].

In our previous study using an MCAo mouse model, we showed that although stroke did not cause detrimental changes to bone microstructure, it prevented the exercise-induced microstructural gains observed in the sham group, even without changes to limb coordination during gait [27]. Since bone and vascular health are tightly coupled, we hypothesized that stroke negatively impacts osteovasculature and inhibits the positive effects of treadmill exercise. In this study, we used the same bones to characterize changes to *in vivo* functional intraosseous perfusion, blood vessel network microstructure, and endothelial cellular composition following ischemic stroke.

## Methods

### Study Design

All procedures were approved by North Carolina State University’s Animal Care and Use Committee. Animals were group housed 4-5 per cage with a 12-hour diurnal light cycle and free access to standard chow and water. Twenty-seven male C57Bl/6J mice (The Jackson Laboratory, Bar Harbor, ME) received either an ischemic stroke (n = 15) or sham surgery (n = 12) at twelve weeks of age (Fig. 1). Mice were housed individually with wetted food and hydrogel packs for four days following surgery during the acute recovery period. After this period, they were returned to their original group cages and split into either daily treadmill exercise groups (n = 6 sham-exercise, n = 8 stroke-exercise) or sedentary control groups (n = 6 sham-sedentary, n = 7 stroke-sedentary). Surgery and exercise groups were randomly assigned to cages. Body mass was measured twice a day for the four days following surgery and weekly thereafter. Stroke recovery and sensorimotor function were assessed daily for four days following stroke and weekly thereafter. Intraosseous perfusion was measured with laser Doppler flowmetry (LDF) during the stroke or sham surgery and then weekly thereafter. After four weeks of stroke recovery with treadmill or sedentary activity, the mice were euthanized with CO2 asphyxiation followed by cervical dislocation. Femora and tibiae from both left (stroke-affected) and right (stroke-unaffected) hindlimbs were collected. A subset of both affected and unaffected femora from six mice (n = 1 sham-sedentary, n = 2 sham-exercise, n = 1 stroke-sedentary, n = 2 stroke-exercise) were fixed for 16 hours in 5% neutral buffered formalin at 4°C and then stored in 1X phosphate buffered saline (PBS) at 4°C until assessment of osteovascular structure. All other bones were fixed for 18 hours in 10% neutral buffered formalin at 4°C and then stored in 70% ethanol at 4°C for assessment of osteovascular composition.

**Figure 1.**
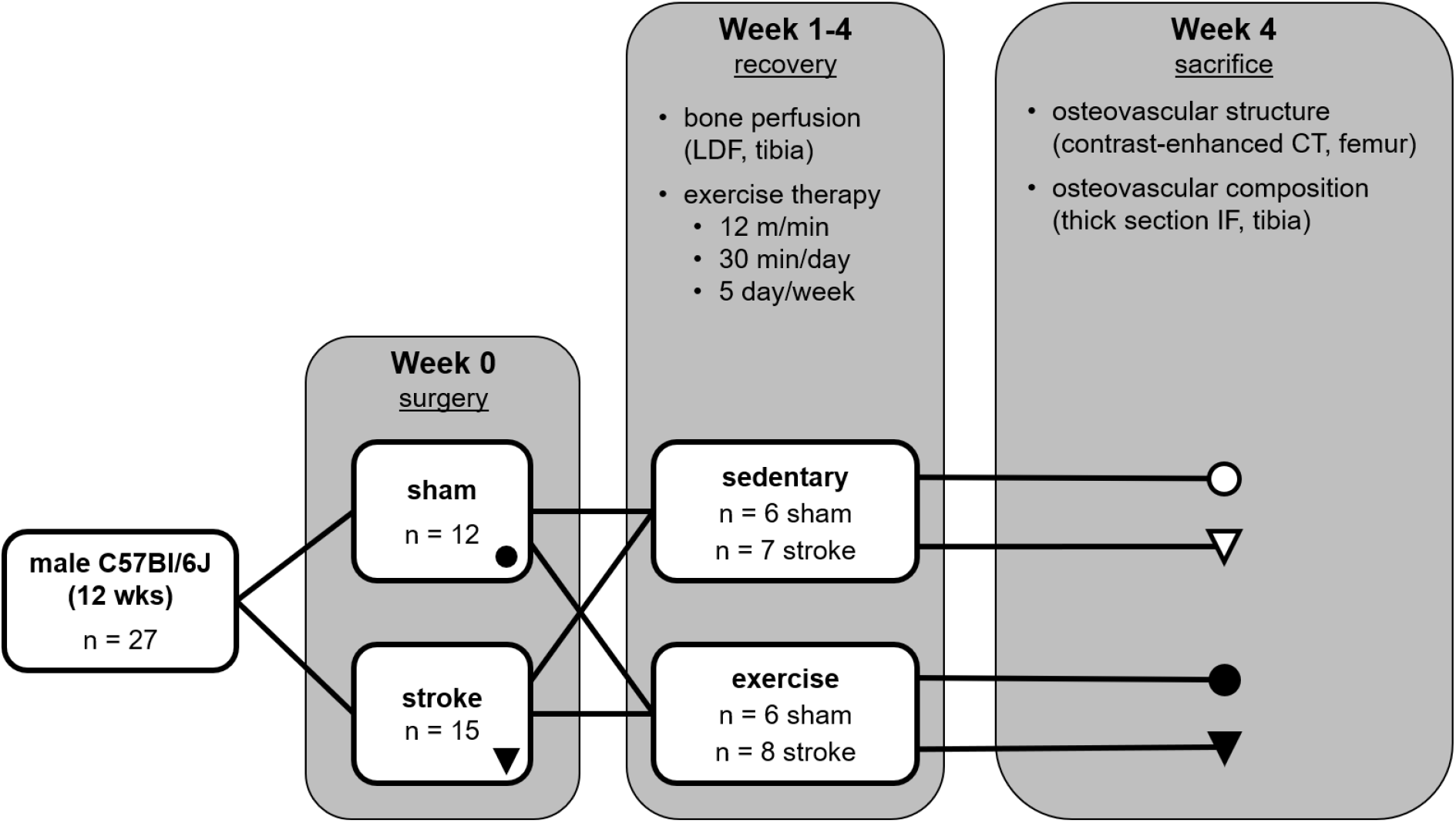
Study design. Mice received stroke or sham surgery, followed by four weeks of treadmill or sedentary activity. Intraosseous perfusion was measured weekly during recovery in affected and unaffected tibiae with laser Doppler flowmetry (LDF). After sacrifice, osteovascular structure and composition were characterized with contract-enhanced computed tomography (CT) and immunofluorescence (IF) microscopy.

### Stroke Procedure

Ischemic stroke was induced with the middle cerebral artery occlusion (MCAo) procedure using aseptic technique [25,26,28]. Anesthesia was induced with 5% isoflurane in a 70:30 N_2_:O_2_ gas mixture, then maintained with about 2% isoflurane. Stroke was induced by occluding the middle cerebral artery with a thin 6-0 nylon monofilament occluder with a silicon-coated tip (Doccol Corporation, Redlands, CA) for 30 minutes. MCAo was monitored using LDF (moor VMS-LDF, Moor Instruments Ltd, Axminster, UK) with a monofilament probe (VP10M200ST, Moor Instruments), ensuring that cerebral blood flow was reduced by 80% relative to baseline and maintained throughout the procedure. The sham procedure was the same as the stroke procedure but without insertion of the occluder. For tibial perfusion, a small, 2-5 mm-long incision was made over the proximal anteromedial side of the left tibia near the metaphysis, avoiding underlying soft tissue and muscle. A small region of the periosteum was scraped away (about 0.5 mm^2^), and a needle probe (VP4 Needle Probe, Moor Instruments) was held firmly against the bone with a micromanipulator (MM3-ALL, World Precision Instruments, Sarasota, FL). The severity of stroke impairments was examined weekly following surgery by assessing sensorimotor function with neuroscores [27].

### Treadmill Exercise

Exercise groups were acclimated to the treadmill (Exer 3/6, Columbus Instruments, Columbus, OH) for two days prior to and two days following (on recovery days 4-5) the stroke or sham procedure. After acclimation, exercise groups performed a standard moderate treadmill exercise regimen (12 m/min, 30 min, 5 days/wk, 5° incline), while sedentary groups were placed on a stationary replica treadmill for a matched time period.

### Bone Perfusion (Tibia)

During the four-week recovery period, intraosseous perfusion was measured weekly in affected (left) and unaffected (right) proximal tibiae using laser Doppler flowmetry (Fig. 2A) [29]. LDF provides a functional measure of blood flow in long bones that is influenced by the amount of blood vessels, blood flow velocity, blood vessel permeability, and blood vessel size [20]. We previously showed that our modified, less invasive LDF procedure can be performed serially without inducing inflammation or gait abnormalities [30]. Using the same method as during the MCAo procedure, an incision was made over one tibia, the periosteum was gently removed, and the LDF needle probe was placed firmly against bone with the micromanipulator. A 30-sec measurement was recorded, the probe was removed and repositioned, and a second 30-sec measurement was recorded. The procedure was repeated for the contralateral tibia. Each weekly perfusion measurement was composed of the weighted mean of the two 30-sec long measurements for that bone.

**Figure 2.**
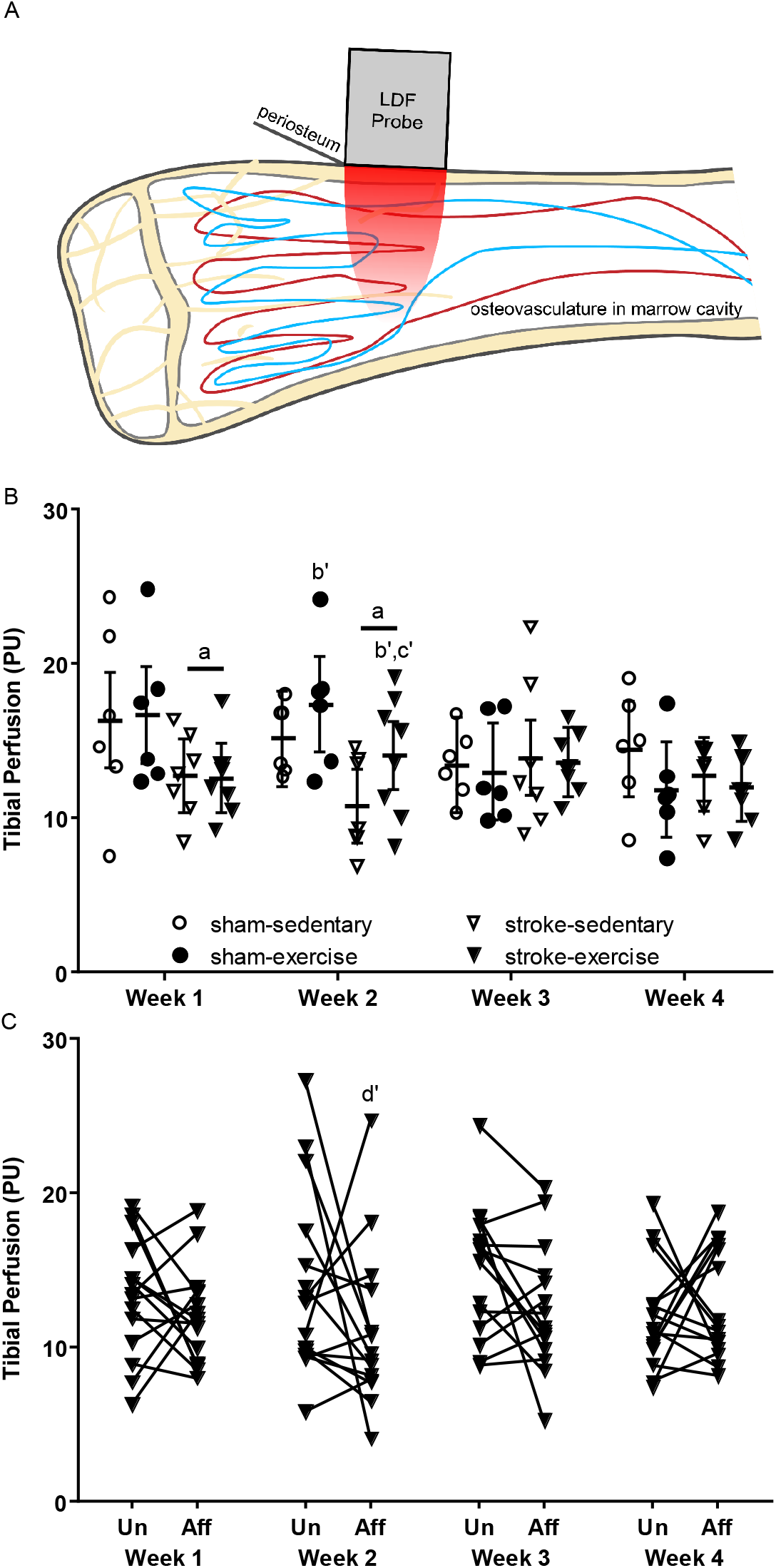
A) Intraosseous perfusion in proximal tibiae measured by laser Doppler flowmetry. B) Perfusion decreased in both limbs at Weeks 1 and 2 post-stroke and was increased with exercise at Week 2 (least squares mean ± 95% confidence interval across both limbs). C) Comparing limbs for stroke, perfusion was nearly decreased in the affected (Aff) relative to unaffected (Un) limb during Week 2. a: p < 0.05 stroke vs. sham (main effect); b’: p < 0.1 exercise vs. sedentary (main effect); c’: p < 0.1 stroke-exercise vs. stroke-sedentary; d’: p < 0.1 Aff vs. Un limb within stroke group.

### Osteovascular Structure (Femur)

Osteovascular structure was examined in the subset of affected and unaffected femora stored in PBS using contrast-enhanced micro-computed tomography (CE-CT). Each femur was cut in half transversely with a scalpel, and the distal halves were incubated in a 3.695 mmol/L staining solution of hafnium-substituted Wells-Dawson polyoxometalate hydrate in 1 mL PBS for 48 hours at 4°C according to a previously described protocol [31]. Samples were scanned with a phoenix nanotom^®^ m (GE Measurement and Control Solutions, Wunstorf, Germany) using 60 kV peak X-ray tube potential, 140 μA X-ray intensity, and 500-ms integration time, and scans were reconstructed at an isotropic voxel size of 2 μm.

Blood vessel and adipocyte microstructure was assessed in a manually selected volume of interest (VOI) that was 1.6-mm long (about 10% average femur length), starting from the distal growth plate and extending proximally into the metaphysis, excluding cortical bone tissue, using the CTAn software (Bruker MicroCT, Kontich, Belgium). Within this VOI, cancellous bone was segmented, binarized via automatic 3D Otsu segmentation, and subtracted from the image, leaving the marrow volume. A manual global thresholding method was used to segment blood vessels and adipocytes. The vessel volume fraction (vessel volume/marrow volume, Ves.V/Ma.V, %), vessel density (Ves.D, #/mm^3^), distribution of vessel thickness (Ves.Th, μm), adipocyte volume fraction (adipocyte volume/marrow volume, Ad.V/Ma.V, %), adipocyte density (Ad.D, #/mm^3^), and distribution of adipocyte thickness (Ad.Th, μm) were quantified using CTAn. Branching parameters of the osteovascular network, including total number of branches, number of junctions, number of triple points (e.g., junctions with three branches), and number of quadruple points (e.g., junctions with four branches), were quantified in BoneJ (FIJI v. 1.51n) [32,33]. The distribution of distances between blood vessel surfaces and bone surfaces (Ves.S-BS distance) was calculated in MATLAB^®^ (R2018, The MathWorks, Natick, MA).

### Osteovascular Composition (Tibia)

Osteovascular composition was examined using immunofluorescence confocal microscopy in bone tissue sections from a subset of affected tibiae (n = 3 sham-sedentary, n = 2 sham-exercise, n = 5 stroke-sedentary, n = 4 stroke-exercise) labeled for markers of non-arterial endothelial cells (EMCN) and endothelial cells in sinuses, arterioles, venules, and capillaries (CD31) [23,34]. Blood vessels that are positive for both EMCN and CD31 (defined as *Type H)* have been shown to regulate bone-vascular crosstalk and couple osteo- and angiogenesis in long bones [24]. Samples were prepared and labeled using a previously described procedure [35], producing 50-μm-thick longitudinal sections of the proximal tibial metaphysis labeled for EMCN and CD31. Nuclear staining was performed with DAPI.

Immediately following staining, sections were imaged at 20X on a Zeiss Laser Scanning Microscope 880 with Airyscan (Carl Zeiss Microscopy, Thornwood, NY), and the areas of CD31-positive, EMCN-positive, and Type H vessels, relative to total area, were quantified in a metaphyseal region of interest (ROI) equal to 10% of the tibia length. CD31-positive regions inside the ROI were manually traced into masks in FIJI, while EMCN-positive regions were automatically binarized with custom code in MATLAB^®^. The CD31 mask, EMCN mask, and the intersection of both masks (Type H) were calculated in FIJI. Osteovascular composition parameters were measured in two sections per bone, and the mean values were used for analysis.

### Statistical Analyses

Statistical models were analyzed with SAS (SAS University Edition v. 9.4, SAS Institute Inc., Cary, NC) with a significance level of 0.05. Statistical models were designed to determine the following: 1) effect of ischemic stroke and exercise on intraosseous perfusion during stroke recovery; 2) acute effect of ischemic stroke on intraosseous perfusion during surgery; 3) effect of stroke and exercise on osteovascular structure and composition; and 4) whether stroke differentially affects intraosseous perfusion or osteovascular structure in the affected vs. unaffected limb.

For analysis #1, LDF measures of perfusion were compared between surgery and activity groups at each timepoint and limb using a mixed hierarchical linear model (procedure GLIMMIX) with interaction between all terms [36]. LDF was measured across four timepoints (Weeks 1-4) on each limb of each mouse. Effect differences between surgery and activity groups were compared within each timepoint (i.e., stroke-exercise vs. stroke-sedentary at Week 2) using least squares means and Tukey-Kramer adjustments for multiple comparisons.

For analysis #2, the stability of the tibial perfusion measurements throughout the surgery was examined using the slope of the 30-minute LDF measurement taken during the surgical procedure. To determine if the slope for each group was different from zero, which would indicate that the perfusion measurement varied and was not stable, an unpaired t-test was performed for the sham group, and a Wilcoxon rank sum test was performed for the stroke group. Slopes were also compared between the stroke and sham groups with a Wilcoxon two-sample test.

For analysis #3, distribution data from CE-CT (vessel thickness, blood vessel surface to bone surface distance) were compared between surgery and activity groups using a mixed hierarchical linear model (procedure GLIMMIX) with interaction between terms. Effect differences were compared between groups within histogram bins (i.e., stroke-exercise vs. stroke-sedentary within bin ‘a’) using least squares means with Tukey-Kramer adjustments for multiple comparisons. All other CE-CT parameters (Ad.V/Ma.V, average Ad.Th, Ad.D, Ves.V/Ma.V, average Ves.Th, Ves.D, branches, junctions, triple points, and quadruple points) were compared between surgery and activity groups using a repeated measures factorial model (procedure MIXED) but without the ‘bin’ repeated factor. Immunofluorescence parameters were compared between surgery and activity groups using a standard two-factor general linear model (procedure GLM) with interaction and Tukey adjustments for multiple comparisons.

For analysis #4, the same repeated measures models from analyses 1 and 3 were used, but least squares means were compared between limbs within surgery group (i.e., affected vs. unaffected side within the stroke group).

Immunofluorescence and LDF data during MCAo are presented as mean ± standard deviation. Data analyzed using least squares means (repeated LDF measures, CE-CT data) are presented as least squares mean ± 95% confidence interval. Repeated LDF and CE-CT data are presented as the least squares mean for both limbs per mouse unless otherwise noted.

## Results

### Bone Perfusion

Tibial perfusion was reduced in both limbs for two weeks following stroke, with 23% lower perfusion in the stroke groups relative to the sham groups at Week 1 (p = 0.0064) and 24% lower perfusion at Week 2 (p = 0.0061), but perfusion levels were similar between surgery groups at Week 3 (p = 0.71) and Week 4 (p = 0.60) (Fig. 2B). Perfusion was nearly greater in exercise groups, relative to sedentary groups, at Week 2 (21%, main effect p = 0.050), primarily driven by the nearly increased perfusion in stroke-exercise (30% relative to stroke-sedentary, p = 0.054), not sham-exercise (relative to sham-sedentary, p = 0.32). Exercise did not affect perfusion in other weeks (p = 0.94 Week 1, p = 0.79 Week 3, p = 0.22 Week 4). Compared to the unaffected tibia, perfusion in the affected tibia was nearly decreased for the stroke group at Week 2 (21%, p = 0.055) but not in Week 1 (p = 0.42), Week 3 (16%, p = 0.11), or Week 4 (p = 0.58) (Fig. 2C). Perfusion was similar between affected and unaffected limbs for the sham group at all timepoints (p = 0.61-0.90).

During the surgeries (occluded for stroke, not occluded for sham), tibial perfusion measurements remained stable throughout, with slopes that were not significantly different from zero for either stroke (p = 0.39) or sham (p = 0.14). The slopes were similar between stroke (−0.0063 ± 0.0415 PU/min) and sham (0.0049 ± 0.0106 PU/min) groups (p = 0.98). These results demonstrate that cerebral ischemia does not directly impact blood supply to the tibia during the occlusion, suggesting that any changes to osteovasculature during the recovery period result from more systemic effects.

### Osteovascular Structure

Blood vessel volume fraction in the distal femur was increased by 38% in the stroke groups relative to sham (p = 0.0061) and decreased by 14% in the exercise groups relative to sedentary (p = 0.031, Fig. 3A-B). Exercise mitigated the effect of stroke, with 17% lower Ves.V/Ma.V in stroke-exercise than in stroke-sedentary (p = 0.027), but did not affect blood vessel volume fraction in sham-exercise compared to sham-sedentary (p = 0.20, Fig. 3B). Neither stroke (p = 0.74) nor exercise (p = 0.24) affected blood vessel density (stroke-exercise: 132.9 ± 215.8 mm^-3^, stroke-sedentary: 62.2 ± 305.2 mm^-3^, sham-exercise: 188.2 ± 215.8 mm^-3^, sham-sedentary: 54.1 ± 305.2 mm^-3^). The vessel network was more branched following stroke, with 26% greater number of branches (p = 0.0089, Fig. 4A), 27% more junctions (p = 0.015, Fig. 4B), 33% more triple points (p = 0.040, Fig. 4C) and 43% more quadruple points (p = 0.017, Fig. 4D), for the stroke group compared to sham. Exercise did not affect any of these network branching parameters.

**Figure 3.**
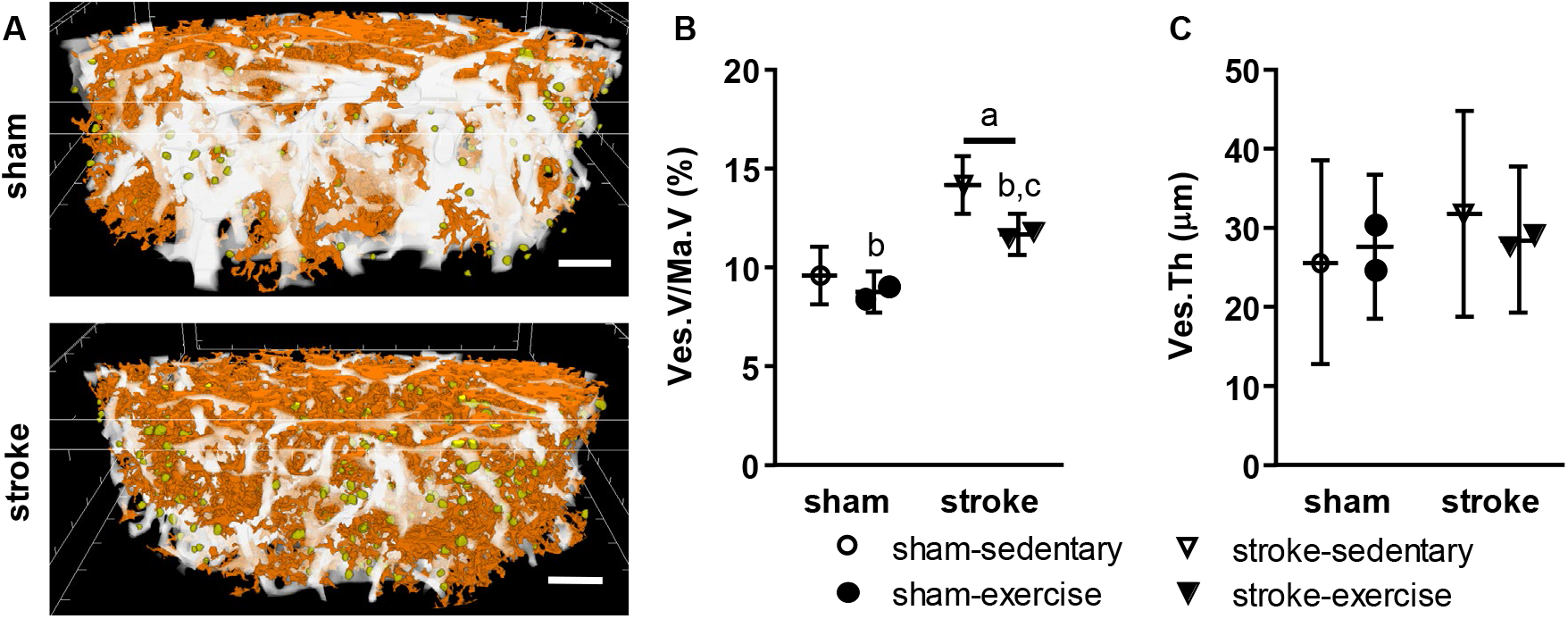
A) Osteovascular structure in the distal femoral metaphysis. B) Blood vessel volume fraction was increased with stroke, while exercise mitigated the effect of stroke. C) Mean vessel thickness was not altered by stroke or exercise. a: p < 0.05 stroke vs. sham (main effect); b: p < 0.05 exercise vs. sedentary (main effect); c: p < 0.05 stroke-exercise vs. stroke-sedentary.

**Figure 4.**
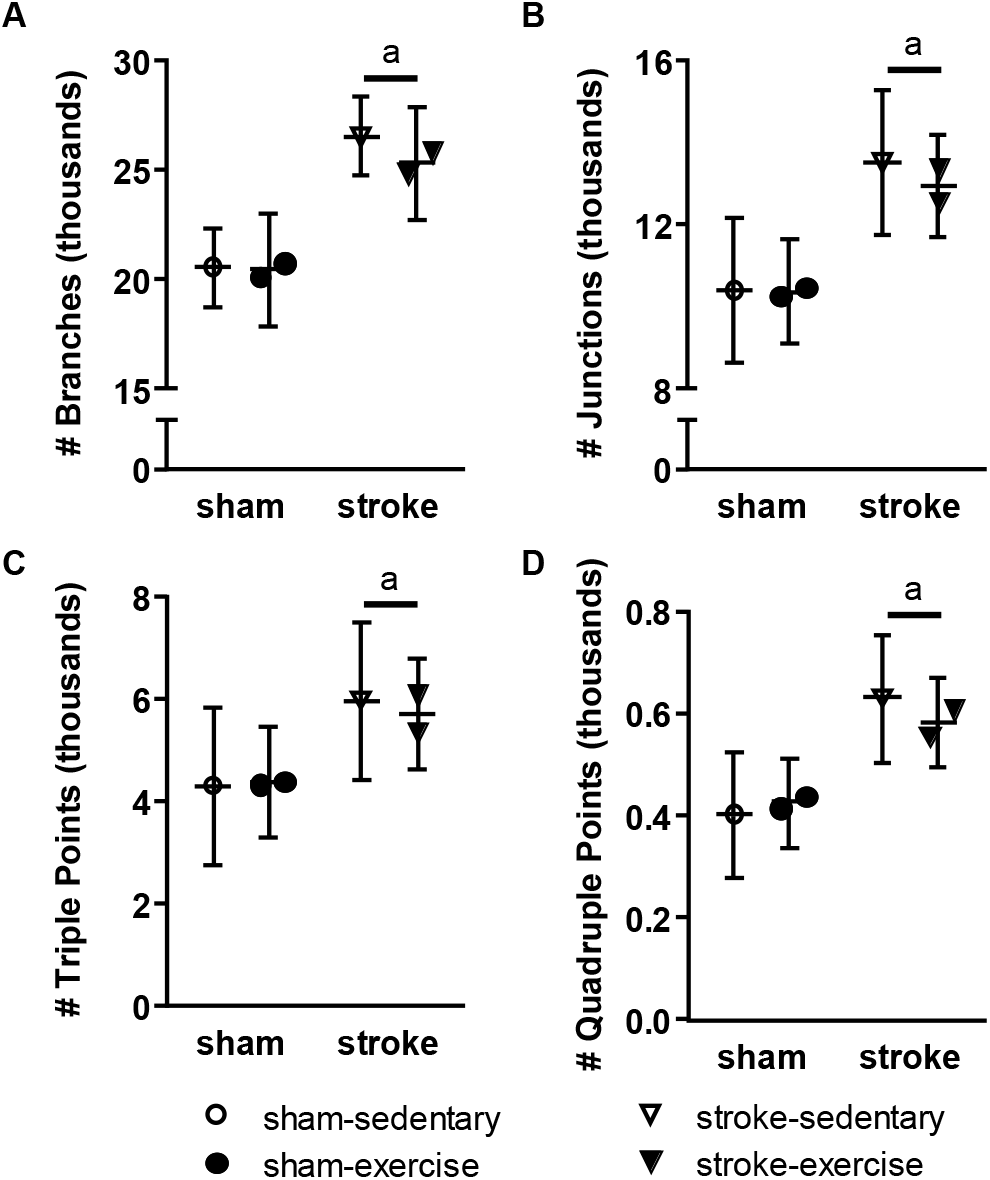
Osteovascular network in the distal femoral metaphysis. A) Vascular branching, B) number of junctions, C) number of triple points, and D) number of quadruple points were increased with stroke. a: p < 0.05 stroke vs. sham (main effect); a’: p < 0.10 stroke vs. sham (main effect).

Neither stroke (p = 0.31) nor exercise (p = 0.84) affected mean blood vessel thickness (Fig. 3C). However, the distribution of Ves.Th was shifted following stroke, with fewer small vessels (6-22 μm thickness, p = 0.029 for 6-10 μm, p = 0.033 for 10-14 μm, p = 0.019 for 14-18 μm, p = 0.056 for 18-22 μm) and more larger vessels (greater than 66 μm thickness, p = 0.010) (Fig. 5A). Exercise mitigated the stroke-induced increase in larger blood vessels (p = 0.0032 stroke-exercise vs. stroke-sedentary), bringing vessel thickness for stroke-exercise to similar values as sham-exercise (p = 0.72) (Fig. 5A). The proximity of blood vessels to bone surfaces was altered by both stroke and exercise, as seen by the shifted distribution of Ves.S-BS distance (Fig. 5B). Compared to sham, the stroke group had 38% fewer blood vessels positioned within 4 μm of bone surfaces (p = 0.041), 19% fewer blood vessels positioned 4-8 μm from bone surfaces (p = 0.0010), and 16% more vessels positioned 52 μm or farther away from bone surfaces (p = 0.036). Exercise had a similar effect as stroke, with 33% fewer blood vessels within 4 μm of bone surfaces (p = 0.017) and 16% more vessels 52 μm or farther away from bone surfaces (p = 0.036) compared to sedentary. The effect of stroke and exercise were driven primarily by changes in the stroke-exercise group, which had significantly fewer vessels positioned within 4 μm and significantly more vessels positioned farther than 52 μm from bone surfaces compared to all other groups (Fig. 5B).

**Figure 5.**
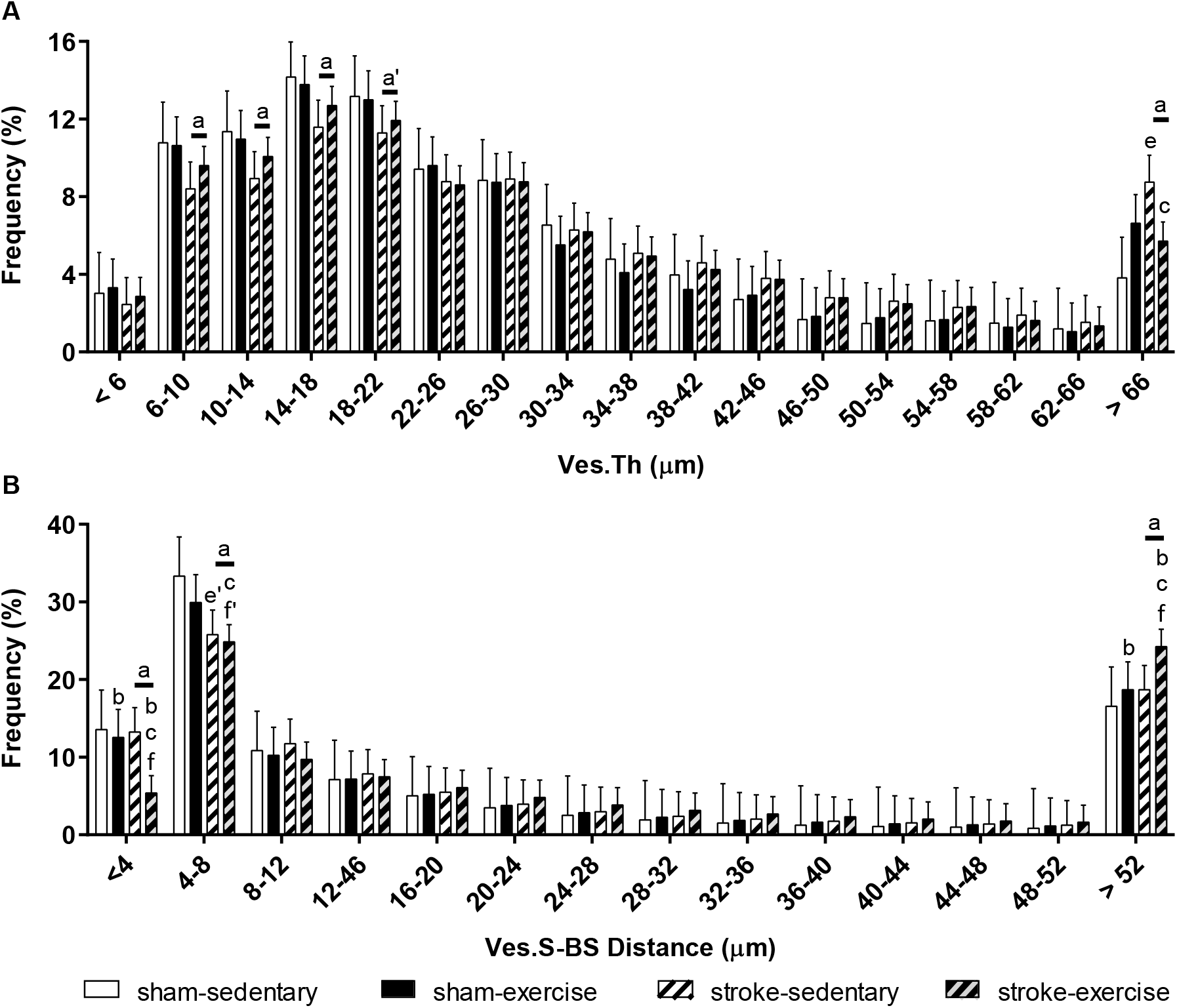
Osteovascular arrangement in the distal femoral metaphysis. A) Distribution of vessel thickness was shifted with stroke, with fewer smaller vessels and more larger vessels. Exercise mitigated the stroke-induced increase in large vessels. B) Both stroke and exercise reduced the relative number of blood vessels nearer to bone surfaces and increased the number farther from bone surfaces, especially for stroke-exercise. a: p < 0.05 stroke vs. sham (main effect); a’: p < 0.10 stroke vs. sham (main effect); b: p < 0.05 exercise vs. sedentary (main effect); c: p < 0.05 stroke-exercise vs. stroke-sedentary; e: p < 0.05 stroke-sedentary vs. sham-sedentary; e’: p < 0.10 stroke-sedentary vs. sham-sedentary; f: p < 0.05 stroke-exercise vs. sham-exercise; f’: p < 0.10 stroke-exercise vs. sham-exercise.

Adipocyte microstructure was not significantly affected by stroke or exercise, with similar values for adipocyte volume fraction, density, and thickness across groups (Table I). However, adipocyte density was nearly increased for stroke-exercise relative to sham-exercise (109%, p = 0.086). Affected and unaffected side measurements were not significantly different in the stroke or sham groups for any of the CE-CT parameters (vessel microstructure, vessel branching, or adipocyte microstructure).

**Table I:** Adipocyte Microstructure in the Distal Femoral Metaphysis

| Trait | Sham |  | Stroke |  |
| --- | --- | --- | --- | --- |
|  | Sedentary | Exercise | Sedentary | Exercise |
| Ad.V/Ma.V (%) | 0.15 ± 0.77 | 0.16 ± 0.55 | 0.15 ± 0.77 | 0.45 ± 0.55 |
| Ad.D (#/mm<sup>3</sup>) | 127.7 ± 166.4 | 113.5 ± 117.7 | 118.4 ± 166.4 | 237.0 ± 117.7* |
| Ad.Th (μm) | 20.5 ± 10.3 | 20.7 ± 7.3 | 20.2 ± 10.3 | 23.1 ± 7.3 |

least squares mean ± 95% confidence interval
\*: p < 0.10 stroke-exercise vs. sham-exercise

### Osteovascular Composition

Neither stroke nor exercise affected the relative area of endomucin-expressing (p = 0.43 and p = 0.54, Fig. 6B) or CD31-expressing (p = 0.18 and p = 0.11, Fig. 6C) blood vessels in the proximal tibial metaphysis of affected limbs. The relative area of Type H vessels expressing both EMCN and CD31 (Fig. 6A) was nearly lower for exercise compared to sedentary (186%, p = 0.059, Fig. 6D) but was not significantly different between stroke and sham (p = 0.20).

**Figure 6.**
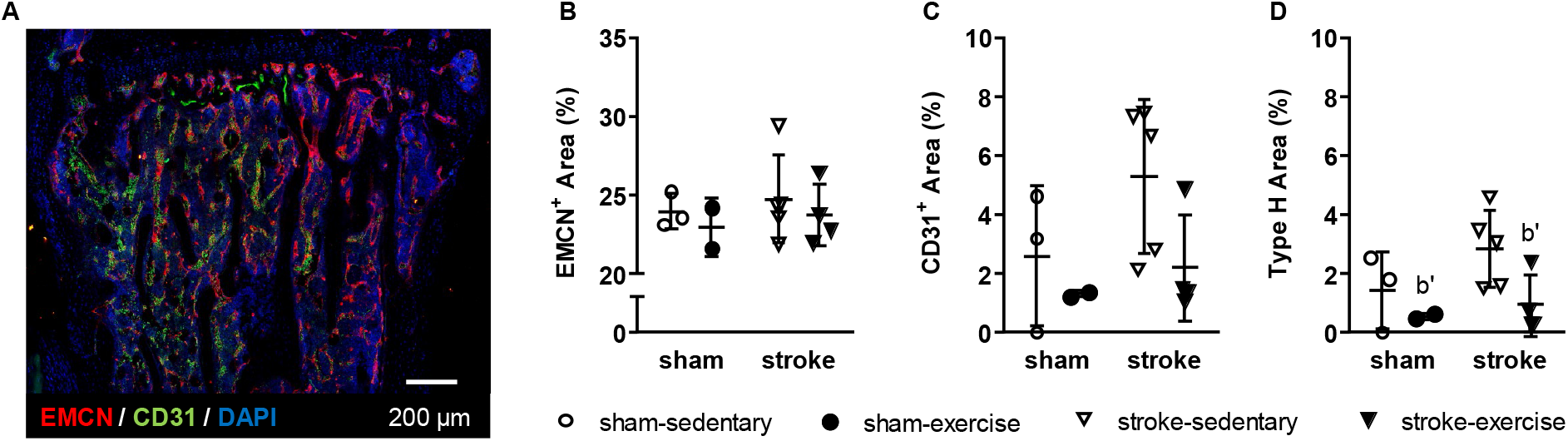
A) Representative longitudinal section in proximal tibia metaphysis, showing osteovascular composition with immunofluorescence. B) Endomucin (EMCN)^+^ and C) CD31^+^ areas were unaffected by stroke or exercise. D) Relative area of Type H endothelial cells (EMCN^+^ and CD31^+^) was nearly reduced by exercise. b’: p < 0.1 exercise vs. sedentary (main effect).

## Discussion

This study is the first to report the effects of ischemic stroke on osteovasculature. Middle cerebral artery occlusion in mice was associated with increased blood vessel volume within the distal femoral metaphysis, primarily due to increased vascular branching and a greater number of larger vessels. These changes followed reduced perfusion in the proximal tibia for two weeks following stroke, with relatively lower blood supply to the affected than unaffected bone in stroke groups but not sham groups. Furthermore, the effects of stroke and exercise on the proximity of vessels to bone surfaces were driven primarily by changes in the stroke-exercise group, yielding fewer blood vessels near to bone surfaces. Because previous studies have shown that active sites of remodeling are associated with increased concentration of capillaries within 50 μm of the remodeling surface compared to non-remodeling bone surfaces [21], these changes to osteovascular structure and function may play a role in bone loss in stroke patients.

Limb disuse via hindlimb suspension has also been shown to decrease bone perfusion in rodents [37,38]. However, in our study, mice remained ambulatory following the mild to moderate severity stroke with no alterations in limb coordination patterns [27], yet bone perfusion was still decreased post-stroke, even in the stroke-exercise group. Together, these results suggest that the osteovascular changes in this study primarily result from more systemic stroke effects rather than from limb disuse. Future work with this induced stroke model may improve understanding about the pathogenesis of skeletal fragility in stroke patients, in particular examining how osteovascular structure and function may contribute.

In this study, stroke decreased functional blood supply within the proximal tibia for two weeks. Perfusion was reduced in both limbs and tended to be lower in the affected side than in the unaffected side during the second and third week of recovery, mimicking the differential decreases in lower leg blood flow observed in previous studies of chronic stroke patients with mild-to-moderate gait asymmetry [17,18]. The decreased perfusion post-stroke was transient, returning to sham levels by the third week of recovery in all stroke groups. Exercise accelerated this recovery, with perfusion in the stroke-exercise group returning to sham-sedentary levels by the second week, indicating that moderate exercise may be a viable strategy for increasing intraosseous blood supply following stroke. Although a number of studies in human stroke patients have shown increased limb perfusion with exercise following stroke [18,39], this study is the first to demonstrate that exercise restores perfusion for the microvasculature within bone, as well.

Perfusion measurements alone do not provide information about changes to the structure or composition of the vascular network, since perfusion could be reduced through decreased vascular density, increased vascular tone, reduced vessel diameter, decreased vascular permeability, or lower blood velocity. Stroke increased osteovascular volume, thickness, and branching in the distal femur by the end of the four-week recovery period. Considering the early transient reductions in bone perfusion in the proximal tibia, the increased osteovascular volume may be a compensatory response to offset perfusion deficits. However, a study examining the progression of these changes over time, and at the same site, is needed to assess the relative timing.

Ischemic stroke is associated with many conditions that could contribute to the reduced intraosseous perfusion observed, including increased vascular tone, increased vascular resistance, or decreased vasodilation. Vascular tone, which constricts blood vessels and decreases perfusion, was higher in cutaneous blood vessels in the paretic hands of patients with ischemic lesions [40] and in denervated arteries in rabbit ears [41], suggesting that stroke-related damage to the central nervous system may contribute to the reduced intraosseous perfusion. Stroke has been associated with increased vascular elasticity (a measure of vascular resistance) in the forearm [15], and resistance arteries in cutaneous blood vessels in the paretic arms of stroke patients were also less responsive to exogenous acetylcholine-induced vasodilation [42], which would increase vascular resistance. However, the effect of stroke on the arteries that supply bone are unknown, and future studies are required to determine whether increased vessel volume and branching are compensatory for changes to vascular tone, resistance, or sensitivity to vasodilators following stroke within long bones. Exercise restored the stroke-related perfusion deficits by the second week of recovery but also reduced osteovascular volume relative to stroke-sedentary. Moderate aerobic endurance training decreased peripheral vascular resistance in rats [43] and increased vasodilator bioavailability in the leg in humans [44], suggesting that the exercise-related perfusion increases may result, at least in part, from improved vascular function. Interestingly, exercise mitigated the stroke-related increases in intraosseous vessel volume, further supporting the idea that the increased vessel volume could be a pathological compensatory response and does not necessarily reflect improved vascular function.

Although stroke increased the amount of blood vessels in bone, the structure and composition of the osteovasculature were also affected, which may contribute to bone loss following stroke. The combination of stroke and exercise decreased the relative number of blood vessels near bone surfaces, which have been shown to be associated with sites of active bone remodeling [21,22,45]. Exercise, in mitigating increased blood vessel volume post-stroke, may have also reduced the number of branches in close contact with bone surfaces. Since exercise did not reduce vessel-bone distance in the sham-exercise group, and since stroke did not decrease vessel-bone distance in the stroke-sedentary group, the combined effect of exercise and stroke may compound the detrimental changes in vessel proximity to bone surfaces. Although the relative area of Type H endothelia was not affected by stroke in the tibia, stroke shifted the distribution of blood vessel diameter in the femur, decreasing the amount of vessels between 6-22 μm in diameter, which matches the size of Type H capillaries known to regulate osteogenesis [23]. Stroke causes chronic inflammation [46], which stimulates angiogenesis [47] and thus may be responsible for the increased vascular volume and branching following stroke, but inflammation also dysregulates Notch signaling [48], the mechanism by which Type H capillaries regulate bone modeling [23,24]. Conversely, exercise nearly lowered the relative area of Type H endothelia in the proximal tibia in both sham and stroke groups but did not affect the number of branches or vessel thickness distribution in the distal femur. The reduction in relative area of Type H endothelial vessels could be due to increased hypoxia-inducible factor (HIF) signaling, which, like Notch, is responsible for Type H activity [24]. Since HIF is released by tissues experiencing hypoxia, HIF in the marrow space may be increased with exercise following stroke if perfusion is negatively affected, and less oxygen and fewer nutrients are able to reach bone tissue. Although some HIF signaling helps recruit osteoblasts [24], compromised bone blood flow restricts oxygen availability to bone cells, stimulating bone resorption [49], and reduced Type H activity could decrease osteogenesis [45]. Despite the increased amount of osteovasculature, the reduction in vessel-bone proximity, resulting primarily from loss of small vessels, may explain the lack of exercise-induced gains in bone microstructure following stroke, which we found in the same region using the same bones as in this study [27].

In this study, we extended previous clinical findings that limb perfusion is reduced following ischemic stroke and demonstrated for the first time, using a mouse model, that intraosseous perfusion is also reduced during early stroke recovery, particularly in the affected limb, and notably even in the absence of limb disuse. These functional deficits occurred in conjunction with changes in osteovascular structure, including potentially compensatory increases in vascular volume, branching, and the number of larger vessels. Further, although exercise mitigated the negative effects of stroke on bone perfusion, the combination of stroke and exercise altered the vessel proximity to bone to a less osteogenic arrangement with fewer small vessels located near bone. These findings suggest that exercise like this moderate treadmill regime may not be beneficial for osteovascular structure, although more studies are needed to examine if these effects extend to other exercise therapies or rehabilitation strategies. Examining potential mechanisms for these changes in osteovascular structure and function following stroke may provide insight for mitigating skeletal fragility in human stroke patients.

## Non-standard Abbreviations and Acronyms

Ad.D: Adipocyte density
Ad.Th: Adipocyte thickness
Ad.V/Ma.V: Adipocyte volume per marrow volume
EMCN: Endomucin
LDF: Laser Doppler flowmetry
Ves.D: Blood vessel density
Ves.Th: Blood vessel thickness
Ves.S-BS: Vessel surface to bone surface distance
Ves.V/Ma.V: Vessel volume per marrow volume

## Acknowledgments

We thank Dr. Eva Johannes and Dr. Mariusz Zareba for confocal microscopy support; and Dr. Consuelo Arellano for statistical consulting. This work was performed in part at the Cellular and Molecular Imaging Facility (CMIF) at North Carolina State University, which is supported by the State of North Carolina and the National Science Foundation.

## SUPPLEMENTAL MATERIAL

### Detailed Methods

#### Stroke Procedure

Following 6-8 hours of fasting, ischemic stroke was induced with the middle cerebral artery occlusion (MCAo) procedure using aseptic technique [1–3]. Anesthesia was induced with 5% isoflurane in a 70:30 N_2_:O_2_ gas mixture, then maintained with about 2% isoflurane. Once mice were anesthetized, fur was shaved over the three incision sites at the anterior neck (for MCAo and sham procedures), the anterior region of the temporalis muscle (for skull LDF probe), and the left proximal tibia (for bone LDF probe). Mice were placed supine on a heated pad, and rectal temperature was maintained at 37°C throughout the procedure (TCAT-2DF, Physitemp Instruments, LLC, Clifton, NJ). LDF was used to monitor both cerebral blood flow (CBF) and perfusion in the proximal tibia using a 785-nm light source (moorVMS-LDF, Moor Instruments Ltd, Axminster, UK). For CBF, a small, 2-5 mm-long incision was made over the right temporal bone, and a monofilament probe (VP10M200ST, Moor Instruments Ltd) was affixed directly to the skull behind the temporalis muscle with cyanoacrylate glue. CBF was recorded with a cutoff frequency of 15 kHz selected for the low-pass filter, along with the automatically applied 20-Hz high-pass filter. For tibial perfusion, a small, 2-5 mm-long incision was made over the proximal anteromedial side of the left tibia near the metaphysis, avoiding underlying soft tissue and muscle. A small region of the periosteum was scraped away (about 0.5 mm^2^), and a needle probe (VP4 Needle Probe, Moor Instruments Ltd) was held firmly against the bone with a micromanipulator (MM3-ALL, World Precision Instruments, Sarasota, FL). Tibial intraosseous perfusion was recorded with a cutoff frequency of 3 kHz selected for the low-pass filter and the automatic 20-Hz high-pass filter.

For the stroke or sham surgery, an incision was made over the neck midline, and the right common carotid artery (CCA), external carotid artery (ECA), and internal carotid artery (ICA) were exposed. In the sham group, saline was added to the neck incision to prevent tissue from drying out, CBF was monitored, and tibial perfusion was recorded for 30 minutes. In the stroke group, temporary ligations were made to the CCA and ICA, while two permanent ligations were made to the ECA, and the vessel was cut between them. Baseline CBF values were collected by loosening the CCA ligation for 2 minutes. The CCA ligation was then retightened, and a thin 6-0 nylon monofilament occluder with a silicon-coated tip (Doccol Corporation, Redlands, CA) was passed through the ECA to the ICA and MCA origin until an 80% reduction in CBF relative to baseline was noted and maintained. The size of the silicon coating was selected based on body mass, per the manufacturer’s instructions, and either a 1-2 or 2-3 mm long, 0.20-0.24 mm diameter coating was selected. Saline was added to the incision to prevent tissue from drying out. The occluding filament was left in place for 30 minutes, and tibial perfusion was recorded. After 30 minutes, the occluder was gently retracted, the ECA was permanently ligated, and temporary ligations were removed. For both sham and stroke procedures, the LDF probes were removed, and an intraincisional injection of bupivacaine (2 mg/kg, Marcaine, Hospira, Lake Forest, IL) was administered to the neck. The neck and skull incisions were sutured closed, and the hindlimb incision was closed with tissue adhesive (VetBond™, 3M Company, St. Paul, MN). Triple antibiotic ointment and 4% lidocaine cream were applied to all incision sites, and a subcutaneous injection of carprofen (7 mg/kg, Rimadyl, Zoetis, Parsippany, NJ) was administered. Bupivacaine was administered via subcutaneous injection for the first two days of recovery. The severity of stroke impairments was examined weekly following surgery by assessing sensorimotor function with neuroscores [4].

#### Bone Perfusion (Tibia)

During the four-week recovery period, intraosseous perfusion was measured weekly in affected (left) and unaffected (right) proximal tibiae using laser Doppler flowmetry (Fig. 2A) [5]. LDF provides a functional measure of blood flow in long bones that is influenced by the amount of blood vessels, blood flow velocity, blood vessel permeability, and blood vessel size [6]. We previously showed that our modified, less invasive LDF procedure can be performed serially without inducing inflammation or gait abnormalities [7]. After 6-8 hours of fasting, anesthesia was induced with 4% isoflurane in pure oxygen and maintained with about 2% isoflurane throughout the 15- to 20-minute-long procedure. The fur over both proximal tibiae was shaved. Mice were placed supine on a heated pad, and the hindlimbs were secured with tape. Using the same methods as described above, an incision was made over one tibia, the periosteum was gently removed, and the LDF needle probe was placed firmly against bone with the micromanipulator. A 30-sec measurement was recorded, the probe was removed and repositioned, and a second 30-sec measurement was recorded. The incision was closed with tissue adhesive, and triple antibiotic ointment and lidocaine cream were applied. The procedure was repeated for the contralateral tibia. Each weekly perfusion measurement was composed of the weighted mean of the two 30-sec long measurements for that bone.

#### Osteovascular Structure (Femur)

Osteovascular structure was examined in the subset of affected and unaffected femora stored in PBS using contrast-enhanced micro-computed tomography (CE-CT). Each femur was cut in half transversely with a scalpel, and the distal halves were incubated in a 3.695 mmol/L staining solution of hafnium-substituted Wells-Dawson polyoxometalate hydrate (K_16_[Hf(α_2_-P_2_W_17_O_61_)_2_]·19H_2_θ) (Hf-POM) in 1 mL PBS for 48 hours at 4°C, with gentle shaking according to a previously described protocol [8]. Samples were scanned with a phoenix nanotom^®^ m (GE Measurement and Control Solutions, Wunstorf, Germany) using 60 kV peak X-ray tube potential, 140 μA X-ray intensity, and 500-ms integration time. A 0.1-mm-thick aluminum filter was applied to reduce beam hardening. Because of the relatively high X-ray attenuation of the Hf-POM-stained samples, a diamond-coated tungsten target was applied. Volumes were reconstructed at an isotropic voxel size of 2 μm using the GE datos|x software with a beam hardening correction setting of 7 and a Gaussian filter setting of 3.

Blood vessel and adipocyte microstructure was assessed in a manually selected volume of interest (VOI) that was 1.6-mm long (about 10% average femur length), starting from the distal growth plate and extending proximally into the metaphysis, excluding cortical bone tissue, using the CTAn software (Bruker MicroCT, Kontich, Belgium). Within this VOI, cancellous bone was segmented, binarized via automatic 3D Otsu segmentation, and subtracted from the image, leaving the marrow volume. A manual global thresholding method was used to segment blood vessels and adipocytes. First, the adipocytes were binarized using opening (sphere with 1-voxel radius), closing (sphere with 3-voxel radius), and two despeckling (remove white and black speckles less than 650 voxels) operations. Then the adipocyte volume fraction (adipocyte volume/marrow volume, Ad.V/Ma.V, %), adipocyte density (Ad.D, #/mm^3^), and distribution of adipocyte thickness (Ad.Th, μm) were quantified. Second, to eliminate edge artifacts due to the partial volume effect, the adipocytes were dilated (sphere with 5-voxel radius) and subtracted from the images, which made the adipocyte edges have similar grayscale values as those of the blood vessels. The images were then segmented to binarize the blood vessels using closing (sphere with 4-voxel radius), despeckling (remove continuous volumes less than 1,050 voxels), closing (sphere with 6-voxel radius), and despeckling (remove continuous volumes less than 5,500 voxels) steps. The vessel volume fraction (vessel volume/marrow volume, Ves.V/Ma.V, %), vessel density (Ves.D, #/mm^3^), and the distribution of vessel thickness (Ves.Th, μm) were calculated.

Branching parameters of the osteovascular network, including total number of branches, number of junctions, number of triple points (e.g., junctions with three branches), and number of quadruple points (e.g., junctions with four branches), were quantified using the ‘Skeleton 3D’ and ‘Analyze Skeleton’ plugins in BoneJ (FIJI v. 1.51n) [9,10], applied to the binarized blood vessels. The distance between blood vessels and bone surfaces was calculated with custom code in MATLAB^®^ (R2018, The MathWorks, Natick, MA). The proximity of blood vessels to bone surfaces was also examined. Similar to the procedure described above, a 1.6-mm-long VOI was manually selected in Slicer (v. 4.11.0) [11], starting at the distal growth plate and extending proximally into the diaphysis, including the cortical bone. The VOI was exported to MATLAB^®^, and bone tissue, adipocytes, and blood vessels were binarized and processed using the same global threshold values and processing steps described above. The distribution of distances between blood vessel surfaces and bone surfaces (Ves.S-BS distance) was calculated using the ‘bwgeodesic’ function in MATLAB^®^.

#### Osteovascular Composition (Tibia)

Osteovascular composition was examined using immunofluorescence confocal microscopy in bone tissue sections from a subset of affected tibiae (n = 3 sham-sedentary, n = 2 sham-exercise, n = 5 stroke-sedentary, n = 4 stroke-exercise) labeled for markers of non-arterial endothelial cells (endomucin, EMCN) and endothelial cells in sinuses, arterioles, venules, and capillaries (CD31) [12,13]. Blood vessels that are positive for both EMCN and CD31 (defined as *Type H)* have been shown to regulate bone-vascular crosstalk and couple osteo- and angiogenesis in long bones [14]. Tibiae were cut transversely at the tibiofibular junction under constant water irrigation with a low speed precision saw fitted with a diamond blade (IsoMet Low Speed Precision Cutter, Buehler, Lake Bluff, IL). The proximal halves of the tibiae were decalcified for 24 hours at 4°C with constant agitation in a solution of 0.342 mol/L ethylenediaminetetraacetic acid (Fisher Scientific, Hampton, NH) diluted in 1X PBS at pH 7.6. Bone samples were then prepared for sectioning and staining with immunofluorescence markers [15]. Briefly, decalcified tissue was infiltrated with a cryoprotectant solution (0.584 mol/L sucrose (S7903, Sigma-Aldrich, St. Louis, MO) and 5.556E-5 mol/L polyvinylpyrrolidone (P5288, Sigma-Aldrich) in 1X PBS) at 4°C for 24 hours with constant agitation. Next, samples were incubated in an embedding media (0.267 mol/L gelatin from porcine skin (G1890, Sigma-Aldrich), 0.584 mol/L sucrose, and 5.556E-5 mol/L polyvinylpyrrolidone in 1 X PBS) at 60°C for 45 min, room temperature for 30 min, and then stored at −80°C until sectioned. In each proximal sample, 50-μm-thick longitudinal sections were obtained using a cryostat at −23°C (HN 525NX, Thermo Fisher Scientific, Waltham, MA). Sections were dried at room temperature for 30 min and then stored at −20°C.

For immunofluorescence staining, slides were equilibrated to room temperature for 30 min, then rehydrated in 1X PBS for 5 min. Sections were permeabilized with a solution of 4.638E-3 mol/L Triton X (T8787, Sigma-Aldrich) diluted in 1X PBS for 20 min at room temperature and blocked in a solution of donkey serum diluted 1:20 in 1X PBS for 30 min at room temperature. Sections were stained using unconjugated antibodies at 1:100 for EMCN (rat anti-mouse sc-65495, Santa Cruz Biotechnology, Santa Cruz, CA), CD31 (hamster anti-mouse ab119341, Abcam, Cambridge, UK), and either vascular endothelial growth factor receptor 2 (VEGFR-2, rabbit antimouse ab2349, Abcam) or osteocalcin (rabbit anti-mouse, ab93876, Abcam), with 7.728E-5 mol/L Triton X diluted in 1X PBS at 4°C overnight. Secondary antibodies were added at 1:200 (goat antirat with AlexaFluor 647 ab150159, Abcam; goat anti-hamster with AlexaFluor 568 A21112 Invitrogen, Carlsbad, CA; and goat anti-rabbit 488 A11034, Invitrogen) with 7.728E-5 mol/L Triton X diluted in 1X PBS for 60 min at room temperature. Nuclear staining was performed with a 7.212E-6 mol/L solution of DAPI diluted in 1X PBS incubated for 10 min at room temperature.

Sections were imaged immediately following staining at 20X on a Zeiss Laser Scanning Microscope 880 with Airyscan (Carl Zeiss Microscopy, Thornwood, NY), and the areas of, CD31-positive, EMCN-positive, and Type H vessels, relative to total area, were quantified. Tile scan images were stitched and processed into maximum intensity projections using the microscope software (Zen 2.3 SP1, Carl Zeiss Microscopy). Regions of interest (ROIs) equal to 10% of tibia length were drawn in FIJI, starting at the proximal growth plate and extending distally into the metaphysis. CD31-positive regions inside the ROI were manually traced into masks in FIJI. Using custom code in MATLAB^®^, EMCN-positive regions were automatically binarized with an adaptive threshold, and then the mask was refined with an opening (disk with 1-pixel radius) followed by a closing (disk with 2-pixel radius) operation. The CD31 mask, EMCN mask, and the intersection of both masks (Type H) were calculated in FIJI. Staining for VEGFR-2 and osteocalcin was indistinguishable from background and could not be quantified. Osteovascular composition parameters were measured in two sections per bone, and the mean values were used for analysis.

#### Statistical Analyses

For analysis #1, LDF measures of perfusion were compared between surgery and activity groups at each timepoint and limb using a mixed hierarchical linear model (procedure GLIMMIX) with interaction between all terms [16]. Each mouse was considered a subject (replicate), assigned randomly to a group comprised of the combination of surgery and activity groups. Surgery groups (sham or stroke) and activity groups (sedentary or exercise) were modeled as fixed factors, limb as a nested within-subject factor, and timepoint as a longitudinal repeated measure. LDF was measured across four timepoints (Weeks 1-4) at each observational unit (each limb of each mouse). Variation among subjects within each group combination (i.e., sham-sedentary) was considered a random effect. Residuals were modeled using a compound symmetry covariance structure. A modified Kenward-Roger approximation was used to calculate denominator degrees of freedom and standard error of fixed effects [16]. Effect differences between surgery and activity groups were compared within each timepoint (i.e., stroke-exercise vs. stroke-sedentary at Week 2) using least squares means and Tukey-Kramer adjustments for multiple comparisons.

For analysis #3, distribution data from CE-CT (vessel thickness, blood vessel surface to bone surface distance) were compared between surgery and activity groups using a mixed hierarchical linear model (procedure GLIMMIX) with interaction between terms. The model was similar to the model used for analysis #1, except the repeated factor ‘week’ was replaced with histogram ‘bins’. Variation among subjects within each group combination (i.e., sham-sedentary) was considered a random effect that was modeled with a heterogenous variance components method across each surgery group (sham or stroke). Variation among each group combination was modeled as an intercept for each group linear predictor. A modified Kenward-Roger approximation was used to calculate denominator degrees of freedom and standard error of fixed effects. Effect differences were compared between groups within histogram bins (i.e., stroke-exercise vs. stroke-sedentary within bin ‘a’) using least squares means with Tukey-Kramer adjustments for multiple comparisons. All other CE-CT parameters (Ad.V/Ma.V, average Ad.Th, Ad.D, Ves.V/Ma.V, average Ves.Th, Ves.D, branches, junctions, triple points, and quadruple points) were compared between surgery and activity groups using a repeated measures factorial model (procedure MIXED) with interaction between terms. Surgery and activity groups were modeled as fixed factors, while limb was modeled as a repeated measure. Residual variance was modeled with compound symmetry covariance, and effect differences were compared between groups using least squares means with Tukey-Kramer adjustments for multiple comparisons. Immunofluorescence parameters were compared between surgery and activity groups using a standard two-factor general linear model (procedure GLM) with interaction and Tukey adjustments for multiple comparisons.

